# fNIRS-Based Neurofeedback for Prefrontal Cortex Modulation: a Proof-of-Concept Study

**DOI:** 10.1101/2025.10.02.679948

**Authors:** Lev Yakovlev, Timofei Ponomarev, Tatiana Gatinskaya, Alexander Kaplan

## Abstract

Functional near-infrared spectroscopy (fNIRS) provides a non-invasive method for monitoring cortical hemodynamics. In this pilot study, we developed a simple neurofeedback system and tested whether participants could volitionally modulate prefrontal cortex blood flow using real-time fNIRS feedback. Preliminary results demonstrate the feasibility of this approach and highlight its potential for future applications in cognitive training and clinical interventions.

## Introduction

Functional near-infrared spectroscopy (fNIRS) is a non-invasive neuroimaging technique that measures relative changes in cortical hemoglobin concentrations, providing insights into brain activity (Ferrari & Quaresima, 2012). Recently, fNIRS has emerged as a promising tool for neurofeedback (NF), offering an alternative to traditional EEG-based methods (Klein et al., 2024). While EEG provides high temporal resolution, fNIRS offers superior spatial resolution, allowing more precise localization of cortical activation. fNIRS-NF has shown positive effects on memory (Hosseini et al., 2016; Hou et al., 2021; Li et al., 2023; Tetsuka et al., 2023), emotional regulation (Li et al., 2025), inhibitory control (Marx et al., 2015; Hudak et al., 2017), and sensorimotor processing (Mihara et al., 2012; Kober et al., 2016). These effects on executive functions and sensorimotor regulation highlight its potential for post-stroke rehabilitation (Mihara et al., 2013; Mihara et al., 2021) and the treatment of autism spectrum disorders (Narita, 2015; Liu et al., 2017), ADHD (Marx et al., 2015; Wu et al., 2022; Barth et al., 2021), anxiety, and depression (Kimmig et al., 2019).

The prefrontal cortex is involved in a wide range of higher-order cognitive functions, including language processing (Price, 2010), emotion regulation (Ray & Zald, 2012), working memory, attention, and inhibitory control (Friedman et al., 2022). Impairments or dysfunction in prefrontal regions have been associated with various pathological conditions (Chini & Hanganu-Opatz, 2021; Pizzagalli & Roberts, 2022) and may also result from normal aging (Korman, 2019). Regardless of their origin, executive functions deficits significantly impact quality of life. fNIRS-based neurofeedback offers a promising approach for both clinical applications and neuroenhancement, with the prefrontal cortex being especially suitable due to reliable signal quality in the hair-free forehead region (Klein et al, 2024).

Recent studies have demonstrated positive effects of fNIRS-based neurofeedback on executive functions. For example, improvement in n-back task performance was reported after several days of training aimed at down-regulating dorsolateral prefrontal cortex (dlPFC) activity (Hosseini et al., 2016). Similarly, significant gains in spatial memory performance were observed in participants who underwent fNIRS-NF training compared to controls (Hou et al., 2021). Consistent findings were reported by Li et al. (2023), where enhanced spatial working memory was associated with increased dlPFC activation. Further, Hudak et al. (2017) demonstrated that dlPFC regulation reduced false alarms in go/no-go tasks and decreased stop-signal reaction time variability in highly impulsive adults, supporting its role in inhibitory control. Comparable improvements were also found in children with ADHD (Marx et al., 2015), highlighting the potential of fNIRS-NF as an intervention for ADHD.

Despite these promising findings, the field of fNIRS-based neurofeedback is still at an early stage, and existing studies remain highly heterogeneous with respect to target brain areas, experimental protocols, optimal trial duration, and number of sessions. Moreover, current approaches often rely on complex laboratory setups, limiting their translation into practical or consumer-level applications. To advance toward real-world use, there is a need for simplified and accessible fNIRS-based systems that still provide valid training of self-regulation.

The aim of this study was to develop a simple fNIRS neurofeedback pipeline that could be implemented in a future consumer-grade device, and to provide initial evidence that such a system enables individuals to self-regulate prefrontal cortical activity. In addition, we tested the hypothesis that a single-session fNIRS-NF training targeting upregulation of the dlPFC would increase mean HbO concentration in the target region.

## Methods

### Participants

Sixteen healthy volunteers (median age=26; 9 female, 7 male) participated in two experimental sessions. All subjects gave their consent to participate in the study.

### Experimental design

All participants completed two experimental sessions, each lasting approximately one hour. The first session involved real neurofeedback (NF), while the second session used sham NF. Session order was randomized across participants, with approximately one week between sessions. During each session, participants sat in front of a computer screen in a laboratory room. Two 3-minute baseline measurements were recorded before and after NF training. Following the first baseline, the NF session began, consisting of 20 trials. Each trial included a 10-second baseline (NF-baseline), a 30-second regulation phase (NF/SHAM), and a 30-second rest period (REST). During the baseline, a fixation cross was presented on the screen. This was followed by the regulation phase, during which a circle appeared and participants were instructed to increase its size and maintain it in green. No explicit strategies were imposed; however, participants were informed prior to the session that cognitive approaches such as recalling words or names, performing mental arithmetic, or visualizing spatial objects could help increase the circle’s size. After the 30-second regulation phase, a small white dot appeared on the screen to indicate the rest period.

### fNIRS neurofeedback

Cortical hemodynamics were recorded using a Brite (Artinis, the Netherlands) continuous-wave fNIRS system. Two wavelengths (752 and 843 nm) were used to estimate changes in oxygenated (HbO) and deoxygenated hemoglobin (HbR). Optodes were placed over the prefrontal cortex (fig.1A), yielding 48 (24+24) measurement channels covering the prefrontal cortex area (dlPFC). The sampling rate was 25Hz. Target channel for neurofeedback training located on F3 position in 10/20 system and corresponded to left dlPFC to upregulate.

**Figure 1.**
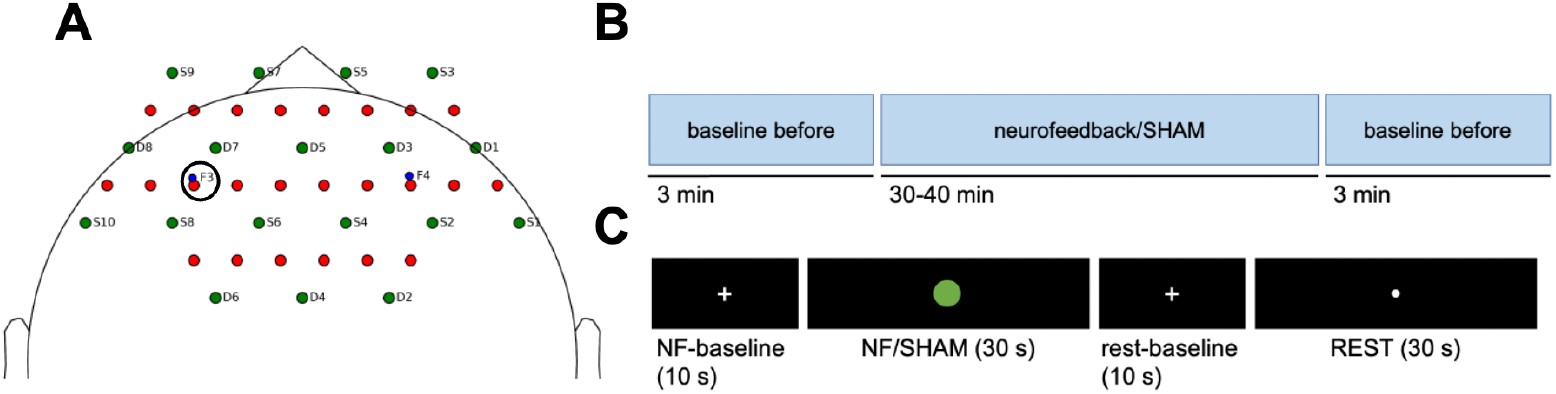
**A**. Optode probe placement. **B**. Experimental session design. **C**. The structure of the trial.

Real-time signal processing and feedback were implemented using a custom Python pipeline, which interfaced with Brite Connect software (Artinis, The Netherlands) via the Lab Streaming Layer (LSL). Raw signals from the target channels were normalized using z-score, and the processed signal was presented to participants as an expanding colored circle, reflecting relative increases or decreases in HbO concentration. To normalize signals for neurofeedback, a 10-second baseline interval immediately preceding each NF trial (NF-baseline) was used. Because the aim of this study was to increase HbO concentration, only the HbO channel was included in the neurofeedback loop.

For each timestamp t in target channel, the normalized signal was calculated as:

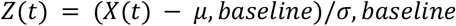

where *X*(*t*) is the current signal timestamp; *µ, baseline* is mean signal value during the 10-second baseline and *σ, baseline* is standard deviation during the same baseline period. During NF trials, the circle’s radius on the screen was scaled according to the z-scored HbO values. Values above z = 5 were displayed in green, indicating strong upregulation, while values below z = -5 were shown in blue, reflecting decreased blood flow. Z-scores between -5 and 5 were represented in yellow, corresponding to baseline levels. In the SHAM-NF session, z-scores for the feedback were calculated randomly.

### Data Analysis

We analyzed the difference in amplitude before and after the whole feedback session and hemodynamics during the session. All following data will be given for our target channel which location corresponds to the F3 EEG electrode in the standard 10-20 system.

fNIRS data were processed using the python MNE package (Gramfort, 2014), including MNE-NIRS toolbox (Luke et al, 2021). Statistical processing was performed using scipy (Virtanen et al., 2020) and pingouin (Vallat, 2018) packages.

The raw light intensity signals were converted to optical density (OD) to mitigate the influence of individual differences in scalp coupling and to improve signal stability. The OD data were converted to hemodynamic changes (millimolar-millimeters, mM·mm) in concentrations of oxygenated and deoxygenated hemoglobin (HbO and HbR) using the Beer-Lambert law (BLL) with a partial pathlength factor (PPF) of 0.1.

For pre- and post-baseline analysis we calculated the mean amplitude for each participant and compared amplitudes using two-way ANOVA (before/after the session neurofeedback/sham feedback during the session) and post-hoc paired T-test. The data for 5 subjects was excluded due to outliers (data that exceeds 3 SD from the group mean). We hypothesized that the neurofeedback session will cause an increase in baseline amplitude.

For NF trials, epochs were extracted for each subject by concatenating data from all experimental runs. For each run, the continuous data were bandpass filtered (0.01 -0.1 Hz) to isolate the hemodynamic response. Events corresponding to the NF, SHAM and corresponding resting state periods were identified. Event codes were standardized across the *NF* and sham conditions. Epochs were defined from -10 s to 30 s relative to the start of the feedback period and baseline-corrected using the pre-stimulus interval.

The further analysis focused on the mean amplitude of the hemodynamic response. For each participant, epochs were averaged separately for each condition. A grand-average waveform was generated by averaging these subject-level averages. Based on the typical temporal profile of the hemodynamic response, the mean amplitude was calculated within a 15–30 s time window for each subject and condition. These mean amplitude values were then entered into a one-way repeated-measures ANOVA to test for a main effect of condition (neurofeedback, sham, 2 corresponding rest conditions). Where the ANOVA revealed a significant effect, post-hoc comparisons were conducted using paired-samples t-tests. The underlying hypothesis was that the neurofeedback condition will show a larger response than rest or sham, thus, a one-tailed test was run.

## Results

### Individual strategies, used by participants

During the feedback sessions, participants employed different mental strategies to increase regional cerebral oxygenation. Although no single unified strategy was prescribed, participants were able to adopt predefined approaches to successfully regulate the circle size. The majority (N = 10) reported that arithmetic operations (e.g., multiplication, division) were particularly effective. As illustrated in Figure 2, the most commonly reported strategies included mental arithmetic, spatial imagery (e.g., navigation or object manipulation), word-based tasks, and planning.

**Figure 2.**
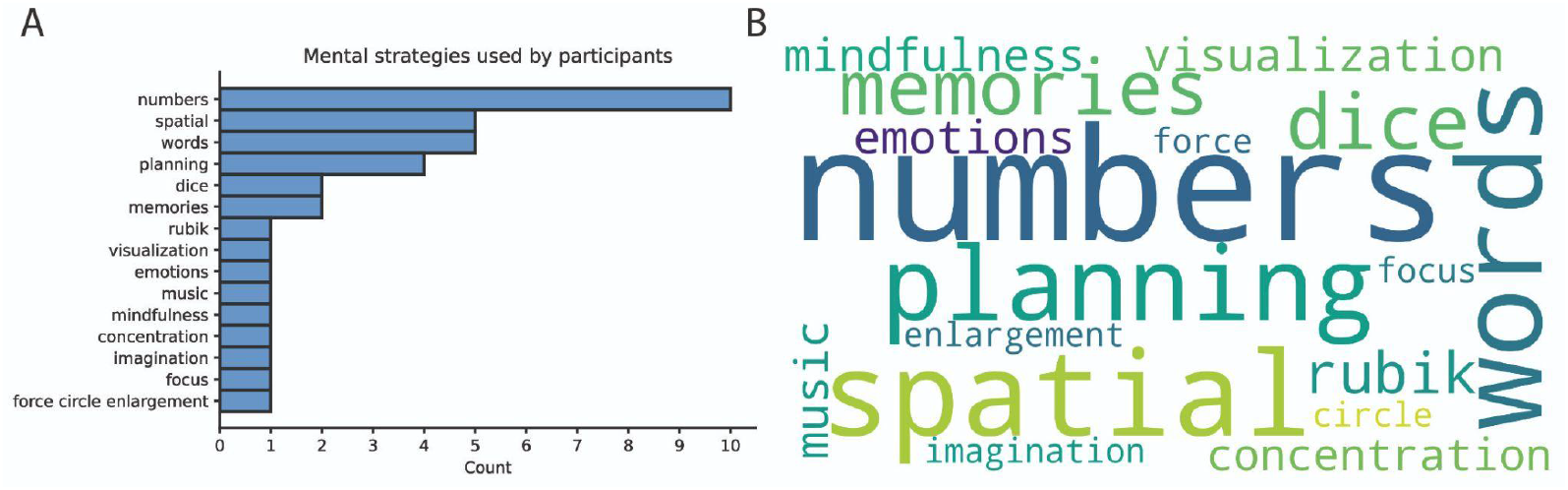
**A**. Histogram of mental strategies used by participants to intentionally increase cortical blood flow. **B**. Word cloud of mental strategies used by participants to intentionally increase cortical blood flow.

### Baseline measurements before and after NF-session

A significant two-way interaction was observed between TIME (pre-vs. post-session) and COND (neurofeedback vs. sham) (*F*_*(1,10)*_*=7*.*29, p=0*.*022, ηg*^*2*^*=0*.*14*). Post-hoc tests indicated a significantly greater amplitude in the post-session baseline for the neurofeedback condition relative to the sham control (*T*_*(10)*_*=-1*.*45, p=0*.*016, d=0*.*55*). Furthermore, a comparison of pre- and post-session baselines within the neurofeedback condition revealed a trend toward an increase in amplitude (*T*_*(10)*_*=2*.*48, p=0*.*089, d=1*.*09*). These results are presented in Figure 3.

**Figure 3.**
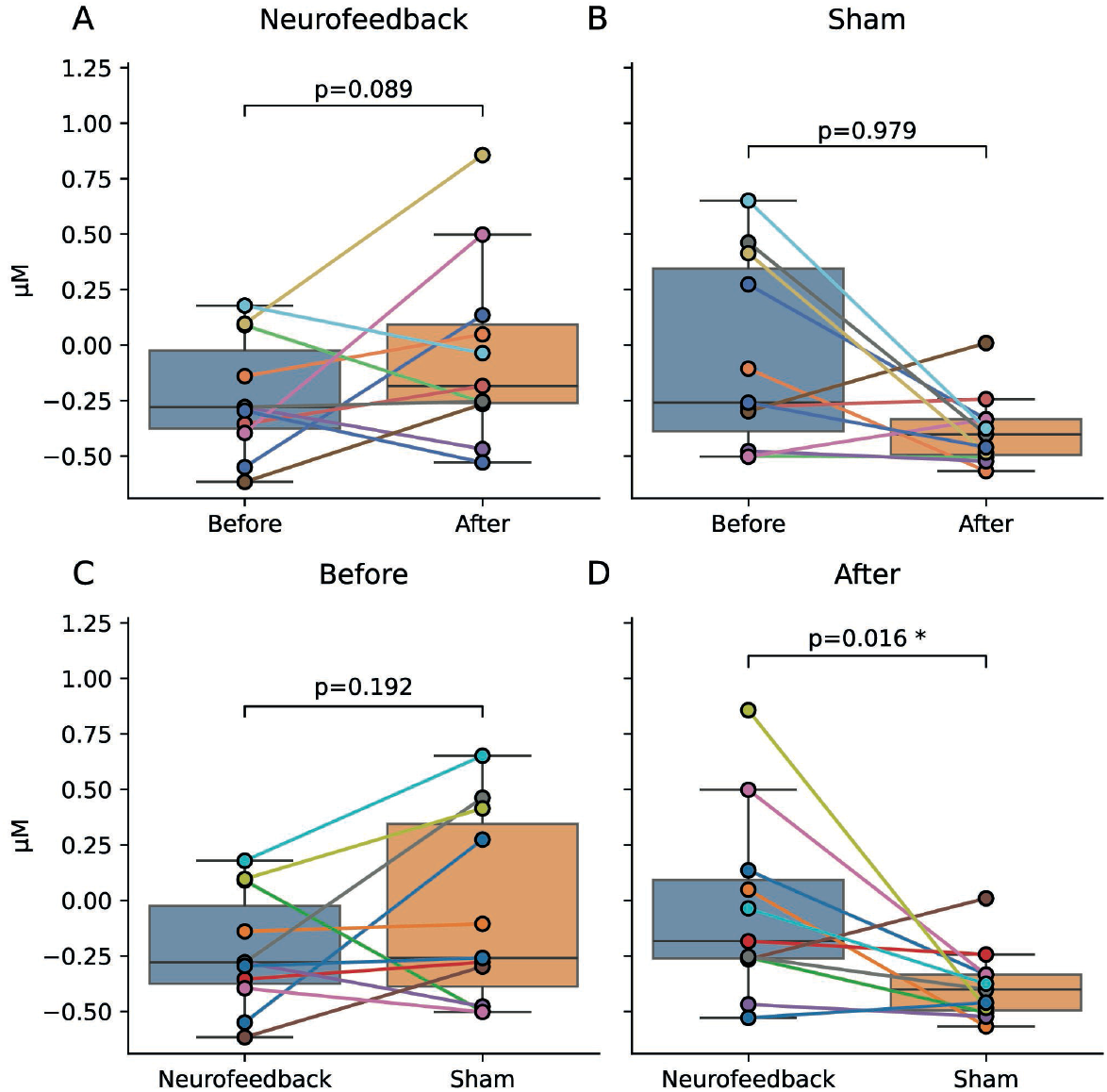
**A**. Mean amplitude before and after a neurofeedback session. **B**. Mean amplitude before and after a sham feedback session. Points and lines represent individual subjects. **C**. Mean amplitude before neurofeedback and sham sessions. **D**. Mean amplitude after neurofeedback and sham sessions. Points and lines represent individual subjects (N=11).

### Neurofeedback session

ANOVA revealed a significant impact of COND on AMPLITUDE at the end of trial periods (*F*_*(3,45)*_*=7*.*29, p=0*.*006, ηg*^*2*^*=0*.*23*). Pairwise comparisons confirmed that HbO concentration in the target channel in the NF-condition is higher than in corresponding rest (*T*_*(15)*_*=2*.*41, p=0*.*015, d=1*.*14*) and sham (*T*_*(15)*_*=1*.*75, p=0*.*050, d=0*.*54*) conditions. Meanwhile, no difference was observed between sham and rest (*T*_*(15)*_*=0*.*83, p=0*.*209, d=0*.*40*). The results are shown in figure 4.

**Figure 4.**
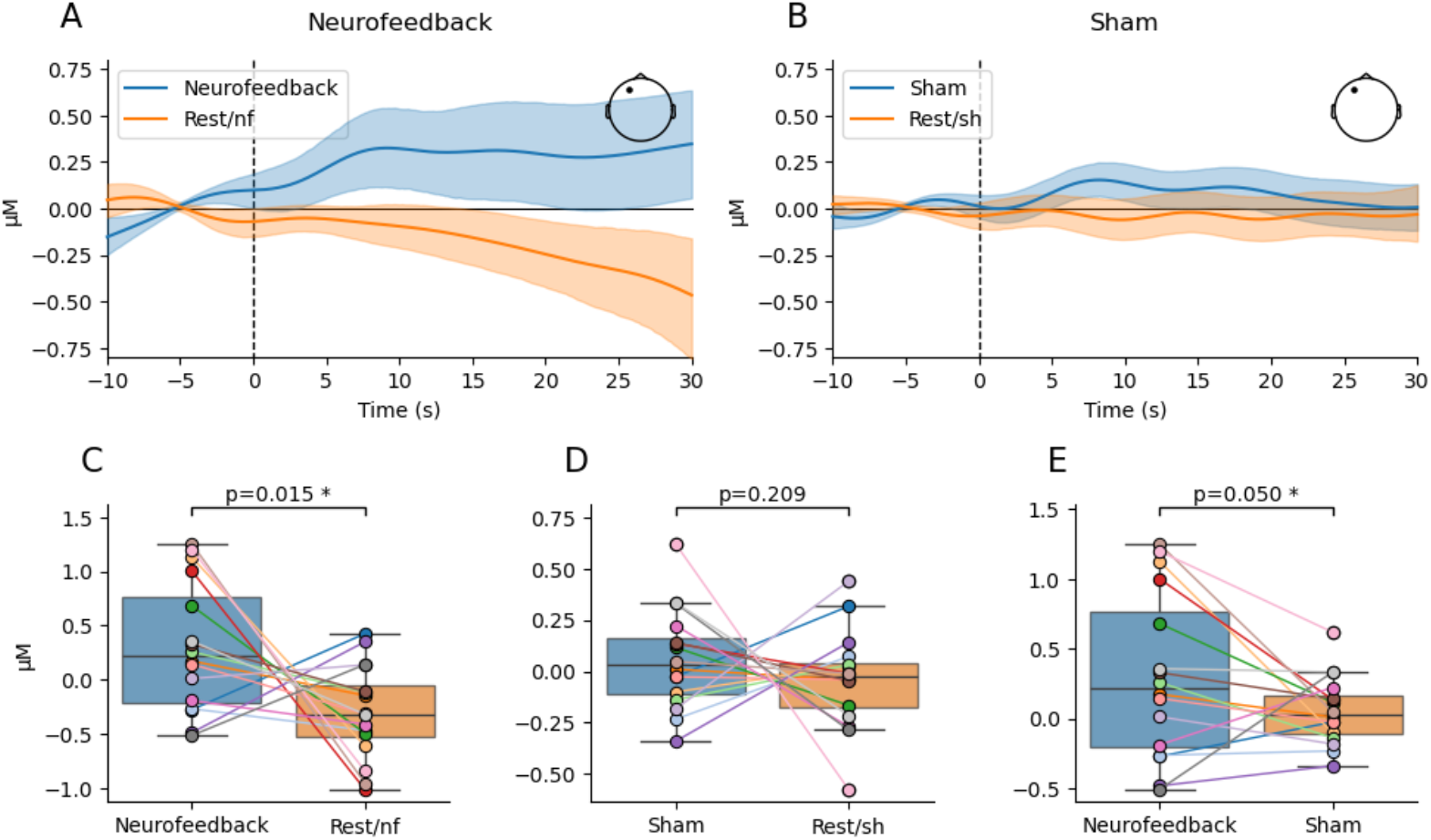
**A**. Mean waveform for the HbO signal at the target position during the neurofeedback session compared to rest. **B**. Mean waveform for the HbO signal at the target position during the sham feedback session compared to rest. **C-E**. Comparisons of amplitude in 15-30s time windows between different conditions. Waveforms presented as a group mean and 95% CI. N=16.

## Discussion

The present pilot study aimed to develop a simple fNIRS-based NF pipeline that could be adapted for consumer-grade applications and to provide initial evidence for its validity in supporting self-regulation of prefrontal cortical activity. The utility of this pipeline was supported by data recorded in sixteen healthy participants. Our results demonstrate that participants in general were able to modulate activity in the left dlPFC during NF training, as reflected by increased HbO concentration in the NF-condition compared to both sham sessions and resting state condition. Importantly, this modulation was observed both during task performance and, to some extent, in baseline activity following training, suggesting short-term effects of self-regulation.

The observed interaction between TIME and COND indicates that fNIRS-based NF can lead to measurable changes in cortical oxygenation beyond momentary fluctuations. Post-hoc tests revealed greater HbO amplitudes after NF-session compared to sham, consistent with our hypothesis that even a single training session may enhance the ability to upregulate prefrontal cortex activity. Although the pre–post comparison within the NF-condition only showed a trend toward significance, the effect size was large, suggesting that a larger sample or extended training could yield more robust effects.

These findings align with previous studies demonstrating the feasibility of fNIRS-NF for modulating prefrontal activity during one single session (Kinoshita et al., 2016; Li et al., 2019).

Our study extends this literature by demonstrating that a minimalistic and technically simple NF setup can elicit measurable changes in cortical activity, representing a critical step toward the development of consumer-level neurofeedback tools. In the study by Kinoshita et al. (2016), a similar single-session approach was used, with both sham and neurofeedback (NF) trials administered within the same session. In contrast, we chose to separate NF and sham sessions on different days. This approach allows participants to focus fully on NF training without potential interference or carryover effects from sham trials. By isolating NF sessions, we aim to enhance learning and consolidation of self-regulation strategies, potentially increasing the effectiveness of each training session.

However, effective and standardized strategies for self-regulating cerebral blood flow still need to be established, and these should remain simple and accessible for participants. Notably, even though participants in our study were provided only with general strategy suggestions (e.g., mental arithmetic, visualization, recalling words), they were still able to increase dlPFC activation, underscoring the feasibility and accessibility of this approach.

At the same time, several limitations should be acknowledged. First, the study employed a small sample size, which limits the generalizability of the findings. Second, the effects observed were short-term; it remains to be tested whether repeated training leads to more stable and behaviorally relevant improvements. Third, only hemodynamic changes were measured, while potential cognitive or behavioral transfer effects (e.g., working memory performance) were not assessed. Future studies should therefore incorporate longitudinal designs, behavioral outcome measures, and larger samples to further validate this approach.

From an applied perspective, these results highlight the potential of fNIRS-NF as the basis for consumer-grade tools aimed at enhancing cognitive performance (Hosseini et al., 2016, Li et al., 2023), supporting mental health (Kimmig et al., 2019), and promoting healthy aging (Trambaiolli et al., 2021). Portable NF devices could be employed for stress regulation, focus training, or executive function enhancement in educational and professional settings. In clinical and preventive contexts, accessible fNIRS-NF may also contribute to rehabilitation after brain injury (Tetsuka et al., 2023), management of ADHD (Marx et al., 2015; Barth et al., 2021; Wu et al., 2022) or autism (Liu et al., 2017), and maintenance of cognitive resilience in aging populations (Trambaiolli et al., 2021). Demonstrating the feasibility of a simple, low-burden pipeline represents a key step toward translating neurofeedback from laboratory-based research into real-world applications.

## Conclusion

Our results provide proof-of-concept evidence that a simplified fNIRS-NF system can enable individuals to self-regulate prefrontal cortical activity. This supports the feasibility of translating fNIRS-NF into practical, user-friendly applications for cognitive enhancement, rehabilitation, and preventive interventions.

## Notes

### Competing Interest Statement

The authors have declared no competing interest.

